# Analysis of prokaryotic genes in an impaired Great Lakes harbour reveals seasonal metabolic shifts and a previously undetected cyanobacterium

**DOI:** 10.1101/2022.12.01.516199

**Authors:** Christine N. Palermo, Roberta R. Fulthorpe, Rosemary Saati, Steven M. Short

## Abstract

As primary drivers of organic matter re-mineralization and trophic carbon transfer, bacteria promote biogeochemical cycling and shape ecosystems globally. We studied bacterial communities spatially and seasonally in an impaired harbour of Lake Ontario by extracting and sequencing community DNA from water samples collected biweekly from different sites. Assembled contigs were annotated at the phylum level, and Cyanobacteria were further characterized at order and species levels. Actinobacteria were most abundant in early summer, while Cyanobacteria were dominant in mid-summer. *Microcystis aeruginosa* and *Limnoraphis robusta* were most abundant throughout the sampling period, expanding the documented diversity of Cyanobacteria in Hamilton Harbour. Functional annotations were performed using the MG-RAST pipeline and SEED database, revealing that genes for photosynthesis, nitrogen metabolism and aromatic compound metabolism varied in relative abundances over the season, while phosphorus metabolism remained consistent, suggesting that these genes remain essential despite fluctuating environmental conditions and community succession. We observed seasonal shifts from anoxygenic to oxygenic phototrophy, and from ammonia assimilation to nitrogen fixation, coupled with decreasing heterotrophic bacteria and increasing cyanobacteria relative abundances. Our data contribute important insights into bacterial taxa and functional potentials in Hamilton Harbour, revealing seasonal and spatial dynamics that can be used to inform ongoing remediation efforts.

## INTRODUCTION

Bacteria facilitate re-mineralization of organic matter and transfer of carbon and nutrients to higher trophic levels, making them essential drivers of global biogeochemical cycles (Falkowski, Fenchel, & Delong, 2008; Fenchel, 2008; Weisse et al., 1990). Microbial ecology has been revolutionized by the advent of metagenomics, which allows the study of entire communities from diverse environmental samples without the limitations of culturing or microscopy-based methods. In addition to diversity and relative abundances, metagenomics offers insights into community functional potential, which has resulted in numerous key discoveries, including those related to the evolution of microbiomes (Hugerth et al., 2015) and global drivers of aquatic microbial community composition (Sunagawa et al., 2015). While comprehensive global surveys of marine microbiomes have been conducted (e.g., Rusch et al., 2007; Sunagawa et al., 2015; Venter et al., 2004), bacterial communities in freshwater environments are relatively understudied.

Here we studied the taxonomic and functional potential of the bacterial communities in Hamilton Harbour, a 21.5-km^2^ embayment located at the western end of Lake Ontario and separated from the lake by a naturally occurring sandbar and the Burlington Shipping Canal (Lawrence et al., 2004). Hamilton Harbour is the largest Canadian port in the Great Lakes and is of major importance to Ontario’s economy (CPCS, 2016). The Hamilton Harbour area has over a century-long history of industrial activity, resulting in a highly polluted harbour containing heavy metals, polychlorinated biphenyls (PCBs), polyaromatic hydrocarbons (PAHs), dioxins, arsenic, and cyanide, among other concerning contaminants (IJC, 1999; Poulton, 1987). Inputs from wastewater treatment plants, sewer overflows, and agricultural and urban runoff have led to high nutrient concentrations, especially phosphorus and nitrogen. Hamilton Harbour is a seasonally eutrophic system that experiences blooms of Cyanobacteria and algae, poor water clarity, and depleted hypolimnetic oxygen concentrations. The harbour was designated an Area of Concern by the International Joint Commission (IJC) and listed as such in the amended 1987 U.S.-Canada Great Lakes Water Quality Agreement (GLWQA). Despite its designation over 30 years ago followed by substantial remediation efforts, it remains one of the most impaired sites in the Great Lakes.

While Hamilton Harbour is in general an extensively studied site, algal diversity and abundance have only been examined during seasonal blooms using microscopic techniques, which can be challenging and subject to errors (Albrecht, Proschold, & Schumann, 2017; Huber et al., 2017; Komarek et al., 2014; Kormas et al., 2011). Furthermore, despite concerns over nutrient and contaminant cycling in the harbour, microbial functional genes have never been analyzed in this system, and therefore the role of the microbial community in cycling and transforming these compounds in this system remains unknown. We aimed to address these knowledge gaps using metagenomics to provide a detailed view of bacterial community diversity, relative abundance, and functional potential during the mid-summer to early fall in Hamilton Harbour.

## MATERIALS AND METHODS

### Sample Collection, DNA Extraction, and Metagenomic Sequencing

Water samples were collected from two sites in Hamilton Harbour as previously described (Palermo et al., 2019). Briefly, 500 ml of water was collected from 1 m below the surface using a Van Dorn bottle sampler from nearshore (43°16’50.0”N 79°52’32.0”W) and mid-harbour (43°17’17.0”N 79°50’23.0”W) locations on July 30^th^, August 13^th^ and 27^th^, and September 10^th^ and 24^th^. Samples were filtered through 0.22 μm pore-size Sterivex capsule filters (EMD Millipore, Burlington, VA, USA), and the filters were stored at −80°C until further analysis. Secchi depth was recorded with each sample, and physiochemical parameters including pH, dissolved oxygen, temperature, redox potential, and chlorophyll a were measured using a YSI 58 sensor (Xylem Inc., Rye Brook, NY, USA). Samples for nutrient analyses were collected from both sites on July 30^th^, September 10^th^, and September 24^th^ following the National Laboratory for Environmental Testing standard protocols. Samples were submitted to the National Laboratory for Environmental Testing (Burlington, Ontario) for sample processing and analysis for the following nutrients: ammonia, chloride, fluoride, sulfate, particulate organic carbon, particulate organic nitrogen, nitrate/nitrite, total dissolved nitrogen, total phosphorus, total dissolved phosphorus, total particulate phosphorus, and soluble reactive phosphorus.

Biomass was recovered from the filters by vortexing for 5 minutes with nuclease free water followed by centrifuging at 6,000 g for 15 minutes. A FastDNA SPIN Kit (MP Biomedicals, Solon, OH, USA) was used to extract community DNA following the manufacturer’s protocol with extra wash steps to maximize removal of environmental contaminants. DNA concentrations were estimated using a NanoDrop ND-1000 UV-Vis Spectrophotometer (NanoDrop Technologies, Wilmington, DE, USA) and sent to MR DNA (Molecular Research LP, Shallowater, TX, USA) for library preparation and sequencing. DNA libraries were prepared using a Nextera DNA Sample Preparation Kit (Illumina, San Diego, CA, USA), and samples were diluted to achieve a recommended concentration of 2.5 ng/μl. For both samples collected on August 27^th^ the maximum sample volume (20 μl) was used for library preparation since the recommended concentration could not be achieved. Libraries were clustered using a cBot System (Illumina, San Diego, CA, USA), and sequenced using 500 cycles on a HiSeq 2500 (2 × 250 bp) system (Illumina, San Diego, CA, USA).

### Metagenome Data Processing

For taxonomic profiles, quality control was performed on raw reads using Sickle version 1.0.0 (Joshi & Fass, 2011) to remove all reads shorter than 50 bp and with Phred quality scores below 30. IDBA-UD version 1.1.3 (Peng et al., 2012) was used to assemble the reads into contigs with a minimum k-mer count of 2, a maximum k value of 20 and a minimum k value of 200. BLASTx searches were conducted using DIAMOND version 0.9.29 (Buchfink, 2015) in frameshift alignment and very sensitive modes. Contigs greater than 500 bp were aligned to the October 2020 NCBI-nr database downloaded from https://ftp.ncbi.nlm.nih.gov/blast/db/FASTA/nr.gz. MEGAN6-LR version 6.14.2 (Huson et al., 2016) was used to assign taxonomic annotations to contigs using the November 2018 MEGAN protein accession mapping file downloaded from: https://software-ab.informatik.uni-tuebingen.de/download/megan6/welcome.html. The Lowest Common Ancestor (LCA) algorithm was used in long read mode with a bit score cut-off value of 150, an e-value cut-off 10^-6^, and a percent identity cut-off value of 70%. To calculate contig relative abundances, reads were mapped back to the contigs using SAMtools version 1.9 (Li et al., 2009) and Bowtie 2 version 2.3.5.1 (Langmead & Salzberg, 2012) in very sensitive and end-to-end modes, as described in Palermo et al. (2019).

For functional potential profiles, raw reads were uploaded to the MG-RAST webserver (Wilke et al., 2016) and processed using default pipeline parameters. Functional potentials of reads and their taxonomic affiliations were annotated using the SEED and RefSeq databases, respectively. Functional annotations were filtered to include only those derived from bacteria and were extracted from MG-RAST for further analysis using an e-value cut-off of 10^-6^, a length cut-off of 35 bp, and a percent identity cut-off of 60%.

### Metagenome Data Analysis

All contigs annotated as bacterial were sorted into one of the following categories based on the NCBI taxonomic classifications: Proteobacteria, Planctomycetes, Actinobacteria, Cyanobacteria, Verrucomicrobia, and other bacteria. The “other bacteria” category contained phyla that were present at less than 5.0% relative abundance, and this category represented less than 8% of the total community in any sample. As they are abundant taxa of concern in Hamilton Harbour, Cyanobacteria were further analyzed at the order and species levels.

There are many tools and databases available to annotate functional attributes of metagenomic datasets. The SEED Subsystems database is a collection of organism-specific functionally related proteins that are manually curated by domain experts, making it more reliable than other functional gene databases such as KEGG (Mao, Zhang, & Xu, 2011). Genes are organized into categories of increasing specificity, with level 1 representing the broadest categorization of functional genes and higher levels increasing in specificity nested within each level 1 category. Descriptions of the categories and the genes they include are available at each level (https://pubseed.theseed.org/?page=SubsystemSelect).

Gene functional affiliations for all samples were averaged at SEED level 1, and relative standard deviations (RSDs) were calculated to illustrate variability between samples in each category. RSDs were calculated by dividing the population standard deviation by the mean and expressing this ratio as a percentage. Several categories were selected for further analysis at SEED level 3, including photosynthesis, nitrogen metabolism, phosphorus metabolism, and metabolism of aromatic compounds, and the breakdown of the subcategories was compared between all samples. Taxa from which functional annotations were derived were profiled for each of the four aforementioned categories. Taxa comprising < 5.0% of the community were summarized in separate supplemental figures.

Pie and bar charts of taxonomic and functional potential profiles were generated in Microsoft Excel. A bubble plot of order-level breakdown of Cyanobacteria was generated using the “ggplot2” package (Wickham, 2009) in RStudio. Heatmaps with sample clustering dendrograms for selected SEED level 3 categories were generated using STAMP version 2.1.3 (Parks et al., 2014). STAMP was also used to produce principal component analysis (PCA) plots of SEED level 1 and photosynthesis level 3 category relative abundances for all 10 samples. Canonical correspondence analyses (CCA) were used to test the relationship between environmental variables and the relative abundances of bacterial phyla, Cyanobacteria orders, and Chroococcales and Oscillatoriales communities using the “vegan” package (Oksanen et al., 2017) in RStudio. Separate CCA models were tested for the entire dataset using tests of 10,000 permutations and the subset of dates for which additional data were available using tests of 719 permutations to compute significance of the model and its variables.

## RESULTS

### Metagenomic Sequencing and Processing Data

The number of reads or contigs at each step in the processing pipeline for bacterial taxonomic profiling are summarized in Table 1. Across all samples, 97.6% to 99.4% of contigs that were assigned a taxonomic label were annotated as bacterial. Although the input DNA for library preparation was lower for both samples collected on August 27^th^, this did not appear to impact the number of reads or contigs generated for these samples. Both samples collected on September 10^th^ had the lowest number of contigs after assembly even though the number of reads post-QC were similar to other samples. With the exception of the September 10^th^ sampling date, all nearshore samples had fewer annotated contigs than the mid-harbour samples collected on the same date.

**Table 1.**
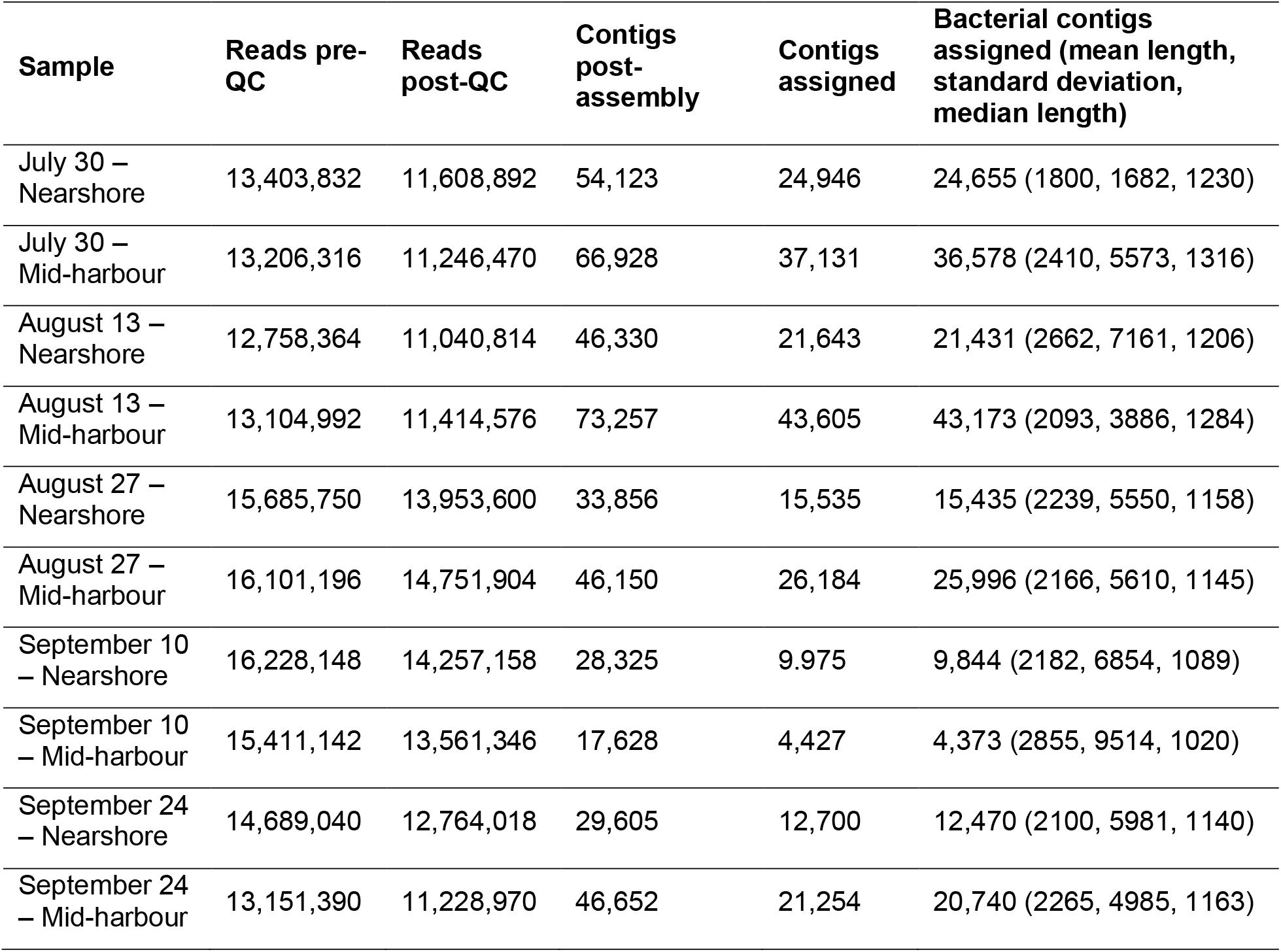
Sequence counts per sample during each step of the taxonomic analysis processing pipeline.

Table 2 summarizes the number of annotated reads in each functional category per sample. For annotations generated using MG-RAST there was a wider range in the number of annotations between samples. However, like the taxonomic annotation pipeline, the samples collected on August 27^th^ had similar contig annotation numbers compared to the rest of the samples, and the samples collected on September 10^th^ contained the lowest number of contig annotations across all samples and almost all gene categories that we explored in detail. Once again, the mid-harbour samples had more annotated contigs than their nearshore counterparts except for the September 10^th^ sampling date.

**Table 2.** Sequence counts per sample from the MG-RAST functional analysis of metagenomes.

| Sample | Bacterial reads assigned | SEED reads assigned | Photosynthesis genes assigned | Nitrogen metabolism genes assigned | Aromatic compound metabolism genes assigned | Phosphorus metabolism genes assigned |
| --- | --- | --- | --- | --- | --- | --- |
| July 30 – Nearshore | 2,845,179 | 1,026,800 | 5,375 | 10,807 | 14,989 | 14,857 |
| July 30 – Mid-harbour | 5,142,341 | 1,850,973 | 17,379 | 19,807 | 25,663 | 27,502 |
| August 13 – Nearshore | 4,119,399 | 1,423,340 | 11,578 | 17,573 | 16,067 | 22,103 |
| August 13 – Mid-harbour | 5,846,566 | 2,030,200 | 26,207 | 25,910 | 21,262 | 30,823 |
| August 27 – Nearshore | 3,807,696 | 1,108,757 | 19,642 | 16,676 | 8,598 | 17,193 |
| August 27 – Mid-harbour | 7,324,859 | 2,087,262 | 34,971 | 31,889 | 15,596 | 32,674 |
| September 10 – Nearshore | 2,173,538 | 627,164 | 8,614 | 8,599 | 6,860 | 8,898 |
| September<br>10 – Mid-harbour | 1,284,769 | 358,345 | 4,731 | 5,000 | 3,896 | 5,310 |
| September<br>24 –<br>Nearshore | 3,219,636 | 867,331 | 14,330 | 13,045 | 9,972 | 13,089 |
| September<br>24 – Mid-harbour | 5,040,321 | 1,450,766 | 21,383 | 21,103 | 16,982 | 22,145 |

### Bacterial Community Composition in Hamilton Harbour

Qualitatively, bacterial relative abundances appeared more similar between sites than between dates (Figure 1). Proteobacteria were highly abundant in July, comprising up to 36.5% to 39.6% of the community at nearshore and mid-harbour sites, respectively. Decreasing at the end of August and early September to below 10% relative abundance, the Proteobacteria increased again in late September to over 30% of the community at both sites. Planctomycetes were low in July and mid-August, comprising less than 5% of the community, but increased at both sites on September 10^th^ to over 25% before decreasing to under 10% relative abundance at the end of September. Actinobacteria were highest in July and early August, decreasing in relative abundance by the end of the sampling period. Changes in the Actinobacteria community over the sampling period were more pronounced at the nearshore site, where they comprised 49.0% of the community in early August, dropping throughout the sampling period until the lowest point of 3.5% relative abundance at the end of September. Actinobacteria at the mid-harbour site followed a similar seasonal trend, though less pronounced, ranging from 9.1% to 21.6% relative abundance. Verrucomicrobia were present at low abundance (< 5.0%) in all samples. Cyanobacteria were present at lower abundances earlier in the season, peaking at the end of September at the nearshore site (54.7%) and at the end of August (64.4%) at the mid-harbour site.

**Figure 1.**
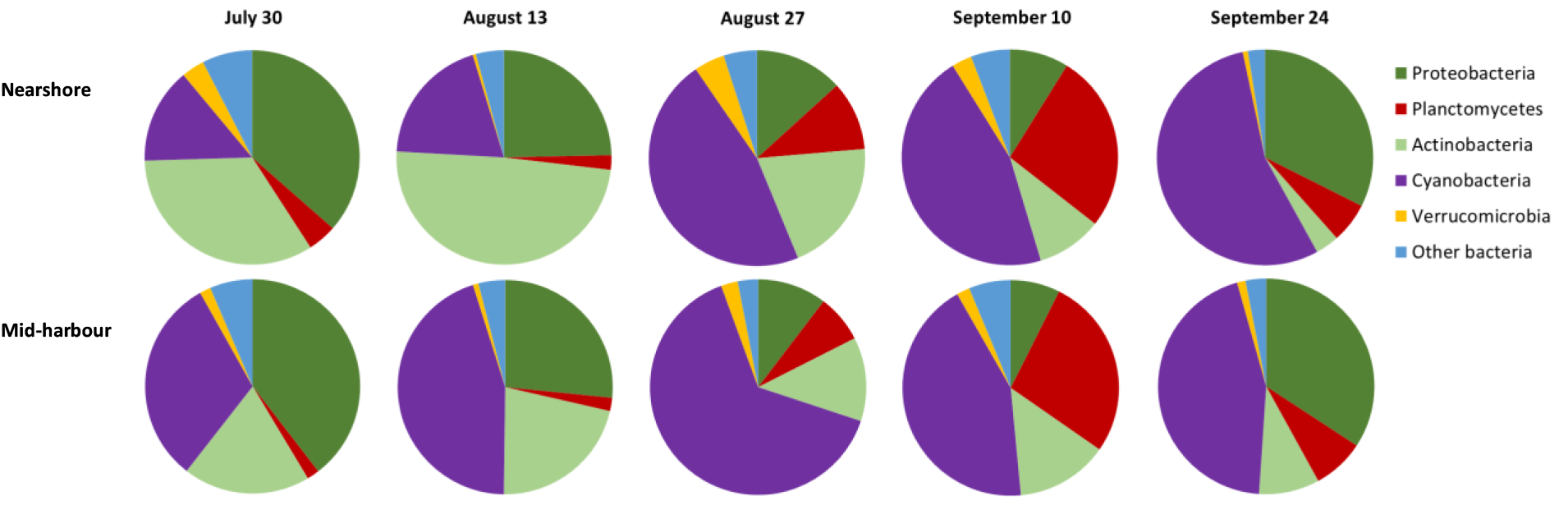
Relative abundances, as a percent of total annotated bacterial contigs, of bacterial phyla detected in each of the ten metagenomes sequenced from Hamilton Harbour.

At the order level, cyanobacterial communities were comprised of primarily of Chroococcales and Oscillatoriales (Figure 2A). Together, these two groups were 83.7% to 92.1% of the cyanobacterial community in each sample. The “Other Cyanobacteria” category contained primarily unclassified Cyanobacteria as well as Pleurocapsales, Chroococcidiopsidales, Spirulinales, and Gloeoemargaritales, most of which were sporadically present in the samples and always less than 1% of the total community. The majority of the Chroococcales community (94.3% to 98.2%) was comprised of unclassified *Microcystis* and *Microcystis aeruginosa* (Figure 2B). The Oscillatoriales community was more diverse, containing several taxa present at low relative abundances, but all samples were dominated by *Limnoraphis robusta* which ranged from 50.3% to 66.2% of the community in each sample (Figure 2C). Synechococcales peaked in July at approximately 5% relative abundance at both sites, dropping to less than 0.4% at its lowest abundance near the end of August. Conversely, Nostocales were lower at the beginning of the sampling period and peaked in mid-September at approximately 6% of the cyanobacterial community at both sites.

**Figure 2.**
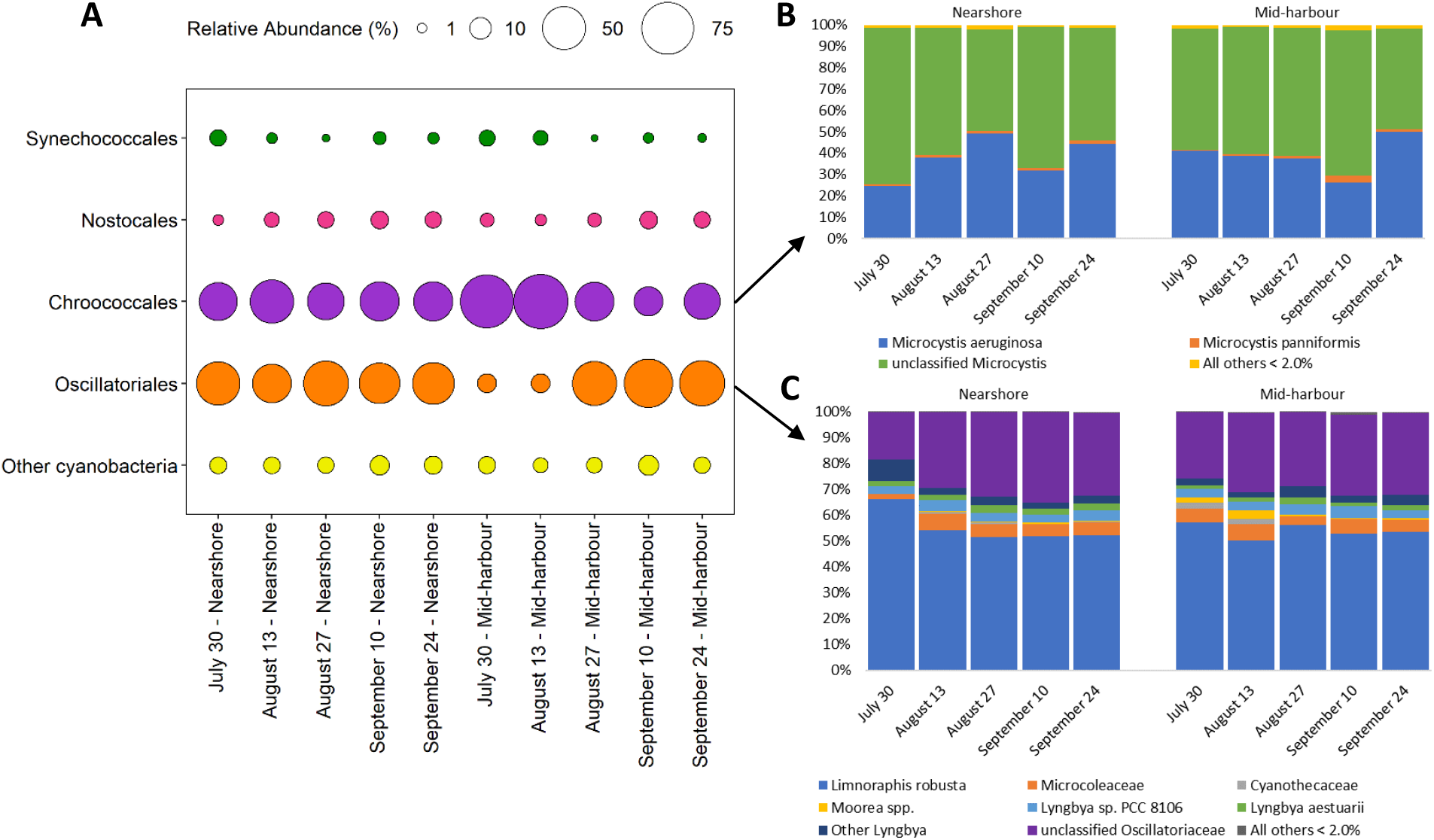
Relative abundances of Cyanobacteria from Hamilton Harbour. A) Bubble plot showing percent relative abundances of cyanobacterial orders accompanied by breakdown of taxa in the largest groups, B) Chroococcales and C) Oscillatoriales. In each panel, the relative abundance shown is the percent of the total number of assigned contigs within each grouping.

### Bacterial Functional Potential

SEED level 1 functional category assignments for all samples are summarized in Figure 3A. The most abundant category was clustering-based systems at 12.8%, while the category with the highest relative standard deviation (RSD) was metabolism of aromatic compounds at 20.3%. Samples collected on the same date clustered closely together in the PCA, and the first and second axes explained 93.7% of the variability in the data (Figure 3B). The following SEED level 1 categories were selected for further analysis: photosynthesis, nitrogen metabolism, phosphorus metabolism, and metabolism of aromatic compounds. As indicated by clustering dendrograms and PCA plot, samples clustered together by date rather than site, indicating a seasonal shift in community functional potential at both sites (Figure 4). In general, the nearshore site showed more pronounced changes over the course of the sampling period than the mid-harbour site.

**Figure 3.**
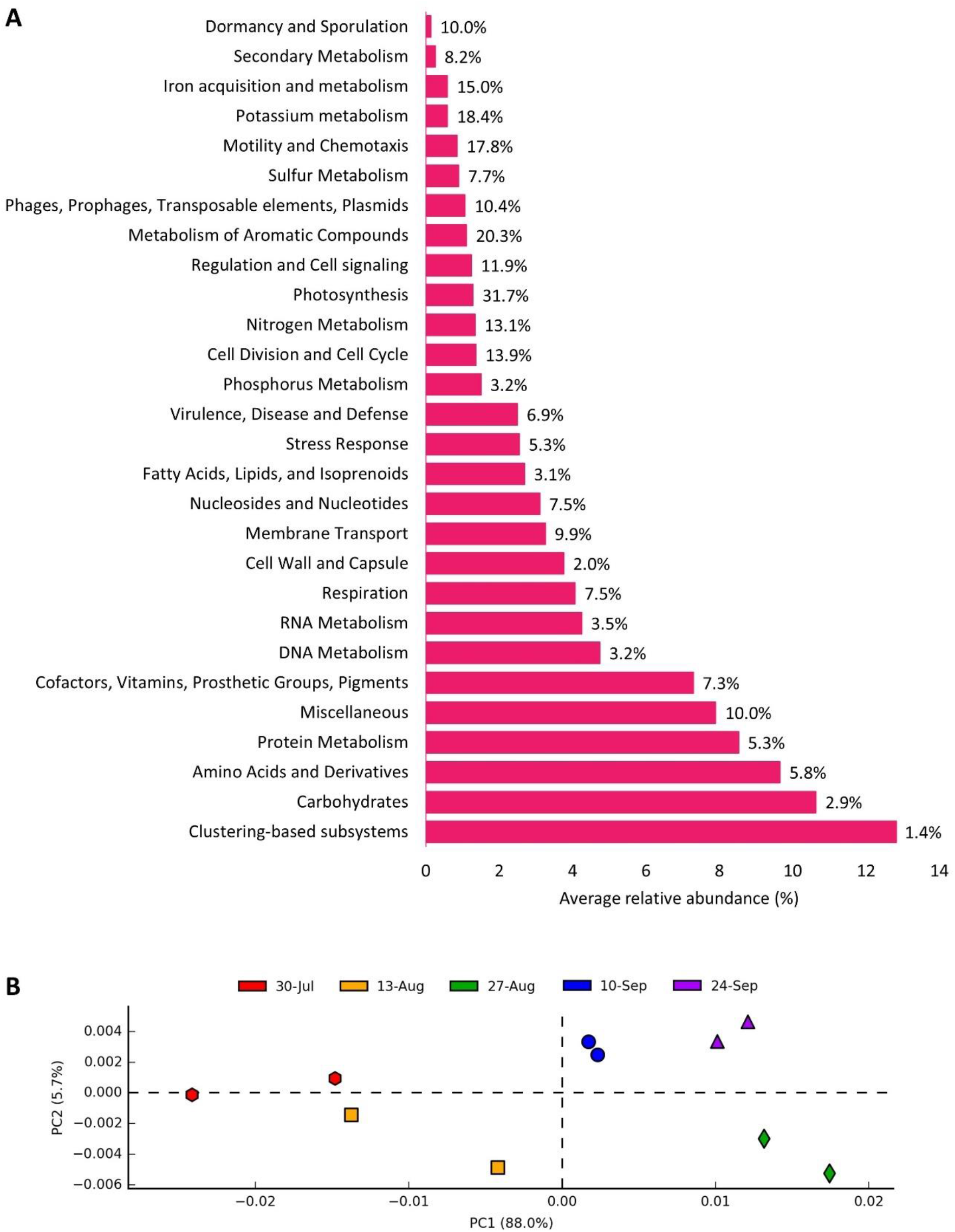
Relative abundances of SEED level 1 categories for assigned bacterial reads. A) Average relative abundances of functional categories for all 10 samples. Relative standard deviation for each category is indicated at the end of the bars. B) PCA of functional category relative abundances by sample collection date.

**Figure 4.**
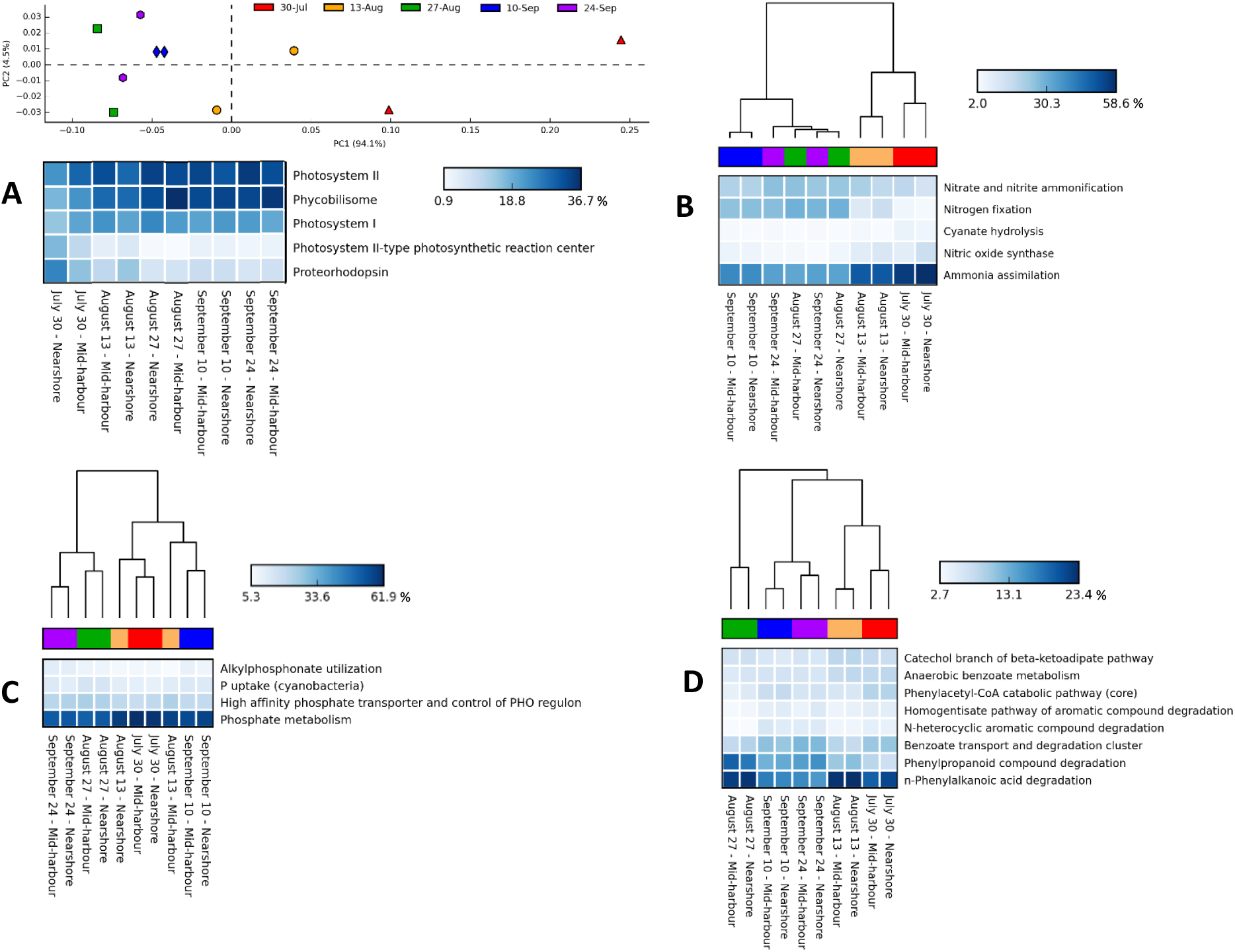
Heatmaps of relative abundances for selected SEED level 3 functional categories below Bray-Curtis dissimilarity dendrograms or PCA for genes encoding A) photosynthesis, B) nitrogen metabolism, C) phosphorus metabolism, and D) aromatic compound metabolism. In each panel, the relative abundance shown is the percent of the total number of assigned reads within each grouping. Only subcategories comprising > 5.0% of the total number genes in each category are included.

Of the genes encoding photosynthetic proteins, subcategories photosystem I, photosystem II and phycobilisome were most abundant overall, and were lowest in July at both sites, increasing later in the sampling period (Figure 4A). Proteorhodopsin and photosynthetic II-type photosynthetic reaction centre genes were highest in July, decreasing later in the sampling period. For genes encoding nitrogen metabolism, the ammonia assimilation subcategory was highest in July and mid-August, decreasing toward the end of the sampling period coupled by an increase in nitrogen fixation (Figure 4B). Nitric oxide synthase and cyanate hydrolysis were also higher in July, decreasing throughout the season, while the nitrate and nitrite ammonification subcategory was lower in July and increased later in the season. Phosphate metabolism was the highest subcategory of all phosphorus metabolism genes, accounting for over 50% of the genes in every sample, with highest relative abundances in July and lowest relative abundances at the end of August (Figure 4C). Relative abundances of genes within the remaining phosphorus metabolism subcategories were relatively consistent across the sampling period and between sites. For metabolism of aromatic compounds, n-phenylalkanoic acid degradation and phenylpropanoid compound degradation were most abundant subcategories, particularly at the end of August, where together they comprised 40.5% and 41.7% of the genetic potential in the nearshore and mid-harbour sites, respectively (Figure 4D). Genes associated with the benzoate transport and degradation cluster peaked early and late in the sampling period, with a decrease in relative abundance on the August sampling dates at both sites.

Abundances of genes within SEED level 1 categories with higher RSDs, such as iron acquisition and metabolism, potassium metabolism, and membrane transport highlight the variability between some categories (Supplementary Figure 1). For example, the potassium metabolism category was dominated by a single gene group; in this case potassium homeostasis was > 78% of the potassium metabolism genes in every sample. Other categories such as membrane transport were comprised of several categories present at lower relative abundances, and no single subcategory was represented at greater than 17% relative abundance in any sample.

### Bacterial Taxa Associated with Functional Categories

In all samples Chroococcales were the most abundant bacterial order associated with photosynthetic genes, ranging from 49.4% to 56.6% at the nearshore site and 45.2% to 71.9% at the mid-harbour site (Figure 5A). Nostocales and Oscillatoriales were both low in July, peaking later in the season, while the opposite trend was observed for Rhizobiales. Other bacteria comprising < 5.0% of the community were also most abundant in July, particularly at the nearshore site, which was also the case for the nitrogen metabolizers.

**Figure 5.**
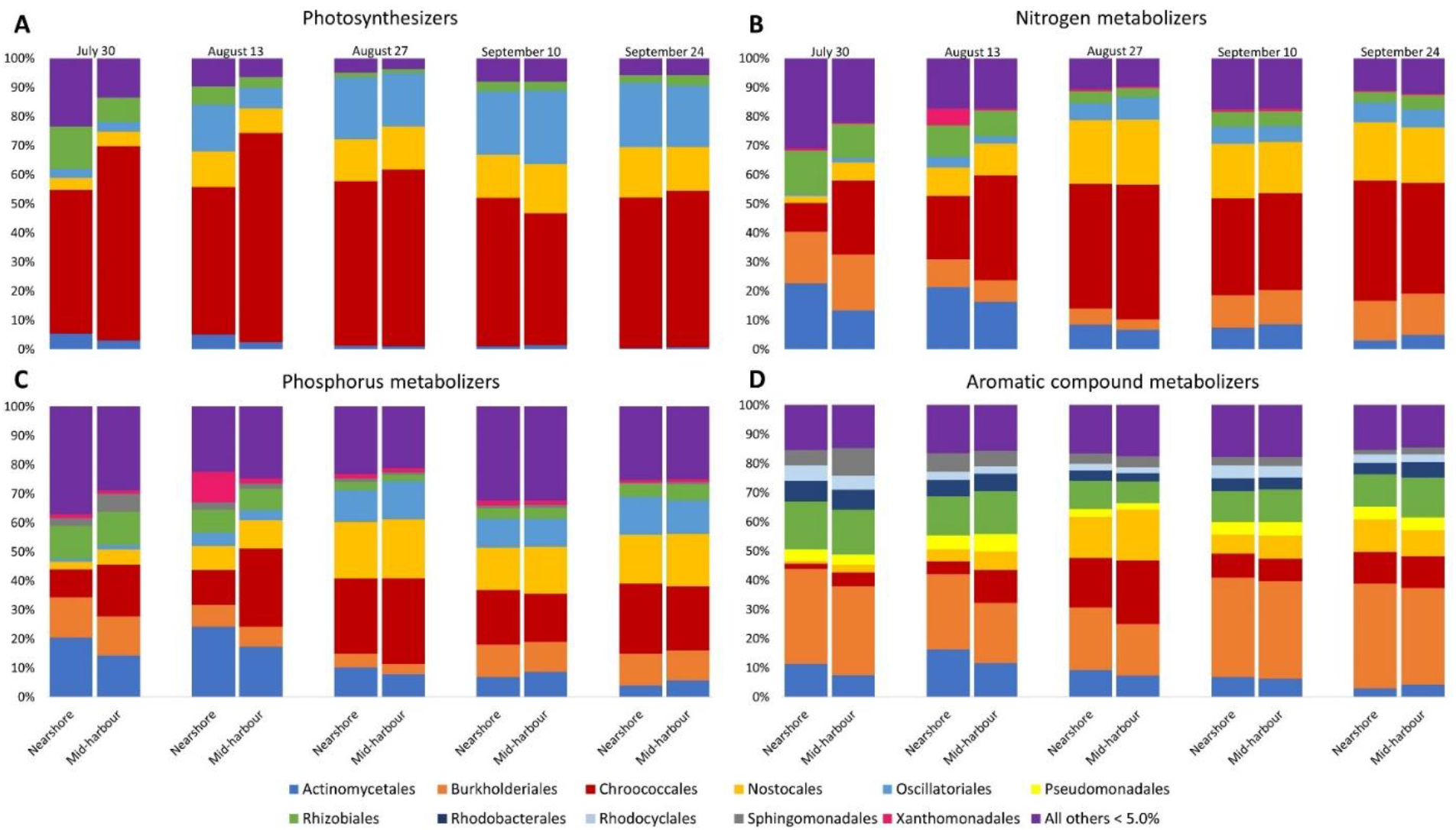
Relative abundances of bacterial orders encoding genes for A) photosynthesis, B) nitrogen metabolism, C) phosphorus metabolism, and D) aromatic compound metabolism. In each panel, the relative abundance shown is the percent of the total number of assigned reads within each grouping. Only bacterial orders comprising > 5.0% of the total number of bacterial orders encoding the genes in each category are included.

Taxa relative abundance trends were similar for bacteria containing genes for nitrogen and phosphorus metabolisms (Figures 5B and 5C). In both cases, Chroococcales and Nostocales were present at low relative abundances in July, peaked at the end of August and remained abundant until the end of September. The opposite trends were observed for Rhizobiales and Burkholderiales, which peaked in July at both sites, and reached their lowest relative abundances at the end of August. Actinomycetales were also higher in July and early August, decreasing over the sampling period to a low point at the end of September. Xanthomonadales were present at low abundances at < 1.0% for nitrogen metabolism genes and < 2.0% for phosphorus metabolism genes, except on August 13 at the nearshore site. In that sample they comprised 5.7% of the bacterial community containing genes for nitrogen metabolism and 10.5% of the bacterial community containing genes for phosphorus metabolism. Of the four categories, phosphorus metabolism contained the highest relative abundances of bacteria comprising < 5.0% of the community, which included 80 unique annotated taxa. The number of bacteria included in this group was similar for nitrogen metabolism (72) aromatic compound metabolism (68), but was much lower for photosynthesis (42).

Burkholderiales were the most abundant bacteria associated with aromatic compound metabolism genes in almost every sample, averaging 28.5% across all samples (Figure 5D). Actinomycetales and Sphingomonadales were higher in July and August, decreasing over the remainder of the sampling period, while Chroococcales and Nostocales were lowest in July, peaking at the end of August, and decreasing again in September. Pseudomonadales, Rhizobiales, Rhodobacterales, and Rhodocyclales had lowest relative abundances at the end of August at both sites but were otherwise relatively stable throughout the sampling period.

### Influence of Environmental Parameters

Canonical correspondence analyses (CCAs) were performed to assess the influence of pH, temperature, redox potential, dissolved oxygen, Secchi depth, and chlorophyll a on the relative abundances of bacterial phyla, Cyanobacteria orders, Oscillatoriales, and Chroococcales in each sample. None of these environmental parameters were significant explanatory variables of changes in bacterial community composition between samples at any of the taxonomic levels examined. For samples collected on July 30^th^, September 10^th^, and September 24^th^, data were available for other environmental parameters including ammonia, chloride, fluoride, sulfate, nitrate/nitrite, total dissolved nitrogen, total phosphorus, total dissolved phosphorus, and soluble reactive phosphorus. Separate CCA modules were generated for the subset of samples for which these additional data were available. These additional parameters were not significant explanatory variables of the changes in bacterial phyla or the Chroococcales communities between samples. The CCA model with total dissolved nitrogen (TDN) as the only predictor variable explained 72.7% (F = 10.66, Pr(>F) = 0.01806) of the variation in the cyanobacterial communities between samples collected on July 30^th^, September 10^th^, and September 24^th^ (data not shown). July 30^th^ had the highest concentration of TDN at both sites, as well as the highest abundances of Actinobacteria and Proteobacteria and the lowest abundances of Cyanobacteria and Planctomycetes. The CCA model explained 44.7% of the variation in the Oscillatoriales communities between the subset of samples (F = 3.23, Pr(>F) = 0.01806). A test of 719 permutations showed that nitrate/nitrite was the only significant environmental parameter (data not shown). Nitrate/nitrite concentrations were highest on July 30^th^ at both sites at approximately 2.3 mg/L, while averaging 1.7 mg/L in all other samples for which data were available. The July 30^th^ samples also had the highest relative abundances of *Limnoraphis robusta*.

## DISCUSSION

In this study we captured the seasonal succession of phototrophic and heterotrophic bacteria over the mid-summer to early fall and observed patterns that were similar across different locations within the harbour indicating that seasonal rather than spatial factors were more important determinants of bacterial community composition. We identified the two dominant cyanobacterial species throughout the sampling period, one of which has never been detected in Hamilton Harbour, and we observed an increase in cyanobacterial genes contributing to photosynthesis as the season progressed. Relative abundances of some gene functional categories like phosphorus metabolism remained stable over the sampling period, while other categories such as nitrogen metabolism were dynamic. In the following discussion, the inclusion of selected gene categories in the analysis and their link to Hamilton Harbour’s history and Area of Concern status will be considered. Given the caveat that some results and conclusions derived from metagenomics are only as good as the databases used for sequence annotation, comparisons between the samples collected for this study revealed fluctuations and patterns in bacterial community diversity, relative abundance, and functional potential.

### Bacterial Communities in Hamilton Harbour

Eutrophication resulting in toxic cyanobacterial blooms has long been a seasonal problem in Hamilton Harbour. The blooms are responsible for Beneficial Use Impairments in the harbour, including degradation of aesthetics and beach closures in the late summer and early fall. Microscopy has been the primary method of studying the diversity and relative abundances of these organisms, but the identification of Cyanobacteria by microscopy can be challenging due to a lack of distinguishable features, species rarity, small cell size, and phenotypic plasticity (Albrecht, Proschold, & Schumann, 2017; Kormas et al., 2011). Moreover, many species cannot be identified, or are likely to be identified incorrectly (e.g., Huber et al., 2017; Komarek et al., 2014; Kormas et al., 2011). Using metagenomics to characterize the cyanobacterial community eliminates many of these challenges and allows us to simultaneously analyze the heterotrophic bacteria community. Despite the potential importance of heterotrophic bacteria on the harbour’s trophic state, the diversity and relative abundance of Hamilton Harbour’s heterotrophic bacterial community has received much less attention than the algal community (Munawar & Fitzpatrick, 2017).

Targeted amplicon sequencing of the 16S rRNA gene has been used to study the bacterial community in Hamilton Harbour (Saati, 2016), showing that the dominant phyla in the summer of 2014 included Actinobacteria, Cyanobacteria, Proteobacteria, and Planctomycetes. Actinobacteria dominated the community in early summer, while Cyanobacteria were the most abundant phyla in mid-summer. Our samples from 2015 showed a similar trend that was particularly pronounced at the nearshore site; higher Actinobacteria abundances were observed early in the sampling period and decreased throughout the season, while Cyanobacteria abundances were lowest in July and increased into September. This complements previous observations from a freshwater reservoir in Spain where early summer communities were dominated by Actinobacteria, which were outcompeted for nutrients by Cyanobacteria later in the summer (Ghai et al., 2014). Our observations of high *Microcystis* spp. abundances, particularly *Microcystis aeruginosa*, were not surprising given longstanding knowledge of their high abundances in Hamilton Harbour during the summer months (Munawar et al., 2017; Munawar & Fitzpatrick, 2018). Interestingly, while *Limnoraphis* spp. have been reported in Hamilton Harbour blooms (Munawar et al., 2017; Munawar & Fitzpatrick, 2018), the most abundant Chroococcales species in our samples, *Limnoraphis robusta*, has not been detected in previous studies. Rather, *Limnoraphis birgei* was the species observed in Hamilton Harbour (Munawar et al., 2017; Munawar & Fitzpatrick, 2018), yet it was not detected in any of our samples. *Limnoraphis robusta* and *Limnoraphis birgei* share similar morphological characteristics (Komarek et al., 2013) and may be challenging to distinguish using microscopic techniques, such as those used in the aforementioned studies. Interestingly, *Limnoraphis robusta* was thought to form blooms only in tropical oligotrophic waters until more recent reports emerged describing bloom formation in eutrophic lakes in North and South America (Komarkova, Montoya, & Komarek, 2016; Kurobe et al., 2013). High relative abundances of *Limnoraphis robusta* in this current study of Hamilton Harbour suggests that this species may be an important component of the mixed algal blooms that occur seasonally in the harbour. If this is the case, it would be the northernmost example of *Limnoraphis robusta* blooms noted in the literature, further expanding the range of conditions under which these blooms form. While eukaryotic algae have been identified as important species in the mixed algal blooms in Hamilton Harbour (Munawar et al., 2017; Munawar & Fitzpatrick, 2018), in our dataset eukaryotes were consistently present at low abundances (< 2%) and were therefore excluded from the analysis.

### Genetic Potential of the Bacterial Community

The top SEED level 1 categories with > 5.0% average relative abundance across all samples were clustering-based subsystems; carbohydrates; amino acids and derivatives; protein metabolism; cofactors, vitamins, prosthetic groups, and pigments; and miscellaneous. These categories contain essential housekeeping genes for cellular maintenance and basic functioning and are typically highly abundant in freshwater environments (e.g., An et al., 2014; Chopyk et al., 2020; Medeiros et al., 2016; Meneghine et al., 2017).

Several functional gene categories were selected for further analysis because of their direct relevance to Hamilton Harbour’s impaired status and with the intent to capture changes in the bacterial community’s metabolic activities during the sampling period. Photosynthesis was selected for more in-depth analysis to gain insight into the cyanobacterial community as Hamilton Harbour is known to support high cyanobacterial populations that occasionally form nuisance blooms during the sampling period. In addition to revealing the relative importance of phototrophy versus photosynthesis in this system, exploring the taxonomic affiliations of photosynthetic functional genes complements the seasonal and spatial taxonomic relative abundance data that was derived using different analysis methods and databases. Nitrogen and especially phosphorus are longstanding nutrients of concern in this seasonally eutrophic system that contribute to the formation of algal blooms and have been subject to seasonal and spatial long-term monitoring in the harbour. The presence of nuisance algae and elevated levels of nitrogen and phosphorus remain *Beneficial Use Impairments* in Hamilton Harbour’s Remedial Action Plan since its inception 30 years ago. The genetic potential and taxonomic affiliations of the bacterial community to uptake and transform these nutrients during the seasonal increase of cyanobacterial abundances could provide key insights into their roles in bloom formation and nutrient limitation in this system. Due to several decades of emissions from industrial and mobile sources, Hamilton Harbour also contains high abundances of aromatic compounds (Sofowote, McCarry, & Marvin, 2008). This category was selected to explore whether the surface water supports a bacterial community with capacity for aromatic compound metabolism that is comparable to unimpacted environments (i.e., primarily contains genes for the metabolism of naturally derived aromatic compounds such as plant or algal secondary metabolites). Alternatively, the functional gene profiles for the metabolism for aromatic compounds may indicate that presence of anthropogenically-derived aromatic compounds is a major influential factor in shaping the community genetic potential. Of these four gene categories selected for more in-depth analysis, photosynthesis, nitrogen metabolism and metabolism of aromatic compounds varied in relative abundances over the sampling period, while phosphorus metabolism remained relatively consistent throughout the season.

As expected, the increase in genes encoding photosystem I, photosystem II and phycobillisome later in the sampling period coincided with an increase in the relative abundance of Cyanobacteria. We observed high abundances of genes encoding proteorhodopsin early in the sampling period, particularly at the nearshore site, which decreased over the sampling period. Proteorhodopsin is a light-activated proton pump that is widely distributed across bacterial genomes in the world’s oceans (Finkel, Beja, & Belkin, 2013; Sieradzki et al., 2018) and is coupled with diverse metabolic pathways (Sabehi et al., 2005). Rhodopsin-based systems were found to be the most abundant light-harvesting systems detected in aquatic metagenomes (Sieradzki et al., 2018), surpassing phototrophy with photochemical reaction centres (Finkel, Beja, & Belkin, 2013). The high abundance of proteorhodopsin genes early in the sampling period may have been influenced by the high diversity of bacteria present at < 5.0% abundance, which comprised over 20% of the bacteria encoding genes associated with photosynthesis in the July 30^th^ nearshore sample. Proteorhodopsin genes are present in a wide variety of bacteria especially Proteobacteria (de la Torre et al., 2003; Dubinsky et al., 2017), which were highly abundant and the dominant group containing these genes at both the nearshore and mid-harbour sites in July. This shift from rhodopsin-based phototrophy to photochemical reaction centre phototrophy as the season progressed was concurrent with the decrease of heterotrophic bacteria and an increase in cyanobacterial relative abundances. Bacteria containing genes for photosynthesis were dominated by a few orders later in the season, compared to earlier in the season when several bacterial orders encoded photosynthetic genes and there was more variation between the two sites.

A similar pattern of a few bacterial orders dominating the functional profiles was also observed in bacteria encoding genes for nitrogen metabolism. The dominant nitrogen metabolism genes shifted from ammonia assimilation to nitrogen fixation over the course of the sampling period, indicating a switch from utilizing nitrogen sources within the system to procuring external sources. As autochthonous nitrogen sources are depleted, the community may undergo a shift to nitrogen fixation, which may have been inhibited by high ammonium levels (Kennedy et al., 1994) earlier in the season. Again, this seasonal shift was reflected in the bacteria encoding genes for nitrogen metabolism, which included a diverse array of heterotrophs and phototrophs in July, developing into a community dominated by Cyanobacteria later in the sampling period. This switch is also reflected in the nutrient data, showing that the highest levels of TDN and nitrate/nitrite were detected at the beginning of the sampling period in July and decreased later in the sampling period.

While there were variations in the relative abundances of groups within the metabolism of aromatic compounds category, the most abundant categories (> 2.0%) are common in a variety of environments (Bai et al., 2013; Puranik et al., 2016; Singh et al., 2018). The most abundant gene categories encoded the metabolisms of n-phenylalkanoic acid, phenylpropanoid and benzoate, all of which have natural sources derived from plants and may have antimicrobial properties (Guven & Onurdag, 2014; Oliveira e Nogueira et al., 2021). The relatively low abundance of the metabolism of aromatic compounds category, averaging 1.1% across all samples, combined with top categories that are common in unpolluted environments, suggests that the primary selective pressures in this category are derived from natural sources.

### Ecological Relevance

Like all temperate inland waters, from July to September, Hamilton Harbour experiences external environmental changes such as decreasing average temperatures and daylight hours. Several nutrients including nitrate/nitrite and total particulate phosphorus (TPP) decreased in concentration from July to September. The PCA plots and Bray-Curtis clustering dendrograms revealed that seasonality rather than location within the harbour was more important for determining bacterial taxonomic and functional potential diversity and relative abundances. Earlier in the sampling period there were greater differences between the nearshore and mid-harbour sites compared to later dates when the two sites were more similar, especially regarding community functional potential. At SEED levels 1 and 3, the relative abundances of genes within the photosynthetic category were more distinct in samples from July and early August compared to samples from late August and September. Later in the season, the communities included a few dominant bacterial orders that accounted for most of the genetic potential of the community.

In environmental microbiomes, genes for essential functions often remain static over time despite changes in microbial diversity and relative abundances (Falkowski, Fenchel, & Delong, 2008). While this tenet was evident for categories such as phosphorus metabolism that remained stable throughout the sampling period, seasonal shifts were evident for categories such as photosynthesis and nitrogen metabolism. The stability of genes associated with phosphorus metabolism reflects the importance of these genes that remained essential despite the various environmental changes and the succession of taxa encoding these genes throughout the season. Conversely, the abundance of genes involved in the metabolism of different nitrogen species changed throughout the season as the community adjusted to changing conditions. This suggests that fluctuating nitrogen conditions exhibit strong selective pressures on microbial taxa and highlights the specialized nature of genes in this category. This is reflected in the CCA of environmental parameters showing that TDN and nitrate/nitrite were significant explanatory variables of the cyanobacterial and Oscillatoriales communities, respectively. Overall, our results demonstrate that several seasonal shifts in prokaryotic taxonomic and functional genes occurred in Hamilton Harbour throughout the summer and early fall.

Here we provide the first report of diversity and relative abundances of the bacterial communities in Hamilton Harbour using metagenomics, revealing seasonal trends and dynamics at different locations within the harbour. We confirmed that *Microcystis aeruginosa* remains a highly abundant nuisance cyanobacterium of concern and identified abundant *Limnoraphis robusta* populations that have not been previously detected in Hamilton Harbour during previous algal bloom monitoring efforts. As the season progressed, we observed an increase of cyanobacterial genes versus heterotrophic bacterial genes contributing to the photosynthesis category, and a shift from proteorhodopsin to photosystems I and II. Similarly, concurrent with increasing relative abundances of cyanobacteria, we captured a shift in nitrogen usage from autochthonous ammonia assimilation to nitrogen fixation. With relevance to ongoing monitoring and remediation efforts in the harbour, changes in relative abundances of certain bacterial groups may indicate the harbour’s nutrient status; for example, the seasonal shift from ammonia assimilation to nitrogen fixation was concurrent with a decrease in the relative abundances of Actinomycetales, Rhizobiales and other heterotrophic bacteria, while Cyanobacteria, especially Chroococcales and Nostocales, increased over the same period.

In closing, the analysis of the functional potential of the bacterial community in Hamilton Harbour presented here revealed that some categories were highly variable (RSD > 10%), reflecting the community’s response to changing resources as the season progressed. Other gene categories were more stable (RSD < 5%) throughout the sampling period, suggesting the presence of generalist gene categories that are essential despite community succession and seasonal changes. Though the fluctuation of genes in the environment allows us to hypothesize about microbial activity, analyzing gene expression (metatranscriptomics) and protein expression (metaproteomics) would allow us to confirm that these genetic changes are physiologically relevant. Future studies would also benefit from a more frequent sampling regime and additional sampling sites to capture more acute responses of bacterial communities to fluctuating environmental conditions.

## ACKNOWLEDGEMENTS

Our thanks to Lewis Molot’s group at York University and Susan Watson’s group at Environment Canada for providing the data on environmental parameters that they collected, measured, and submitted for processing to the National Laboratory for Environmental Testing. We are grateful to the many Environment Canada and Climate Change Technical Operations boat crews for their assistance.

## AUTHOR CONTIBUTIONS

Conceptualization, C.N.P., R.R.F., R.S. and S.M.S.; Methodology, C.N.P., R.R.F., R.S. and S.M.S.; Validation, C.N.P. and S.M.S.; Formal Analysis, C.N.P. and S.M.S.; Investigation, C.N.P. and S.M.S.; Resources, R.R.F. and S.M.S.; Data Curation, C.N.P. and S.M.S.; Writing – Original Draft Preparation, C.N.P. and S.M.S.; Writing – Review & Editing, C.N.P., R.R.F., R.S. and S.M.S.; Visualization, C.N.P. and S.M.S.; Supervision, R.R.F. and S.M.S.; Project Administration, R.R.F. and S.M.S.; Funding Acquisition, R.R.F. and S.M.S.

## DATA AVAILABILITY STATEMENT

Data generated during this study are available in the MG-RAST repository, under the following accession numbers: mgm4683865.3, mgm4683866.3, mgm4683867.3, mgm4683868.3, mgm4683869.3, mgm4683870.3, mgm4683871.3, mgm4683872.3, mgm4683873.3, mgm4683874.3.

