## Supplementary Figure 1 for "Analysis of prokaryotic genes in an impaired Great Lakes harbour reveals seasonal metabolic shifts and a previously undetected cyanobacterium"

A

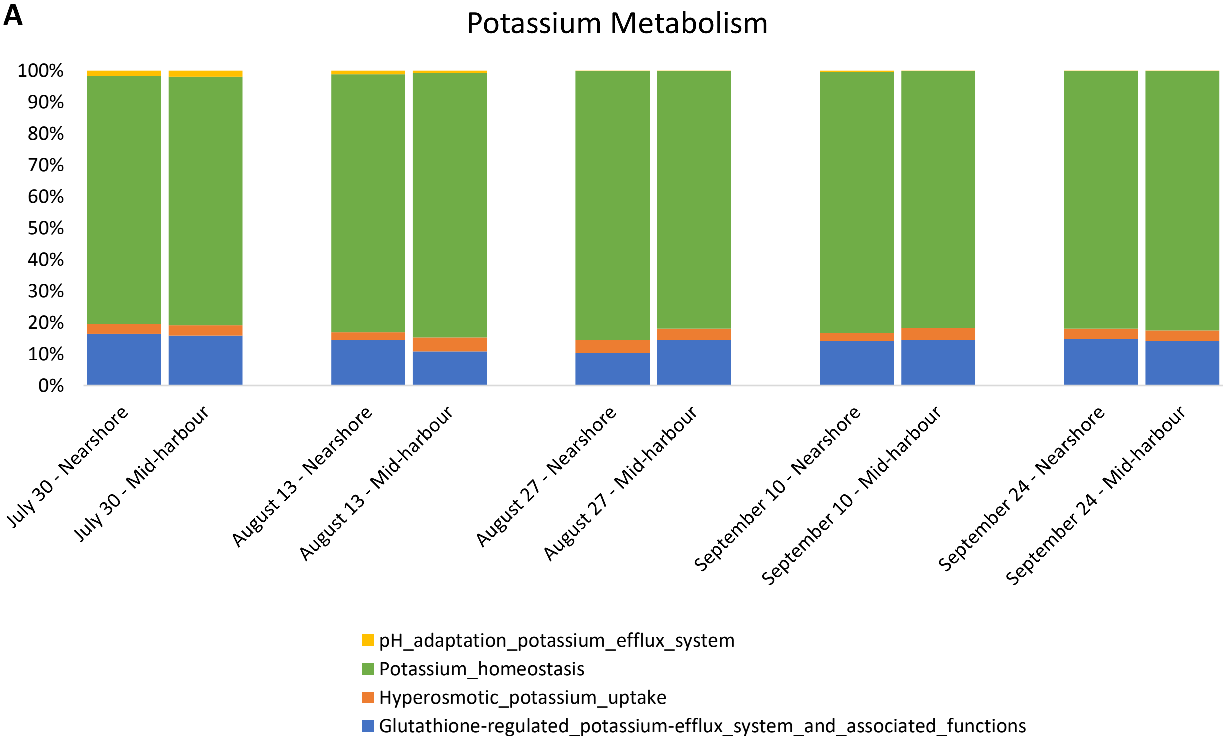
Potassium Metabolism
100%
90%
80%
70%
60%
50%
40%
30%
20%
10%
0%
July 30 - Nearshore
July 30 - Mid-harbour
August 13 - Nearshore
August 13 - Mid-harbour
August 27 - Nearshore
August 27 - Mid-harbour
September 10 - Nearshore
September 10 - Mid-harbour
September 24 - Nearshore
September 24 - Mid-harbour
pH_adaptation_potassium_efflux_system
Potassium_homeostasis
Hyperosmotic_potassium_uptake
Glutathione-regulated_potassium-efflux_system_and_associated_functions

B

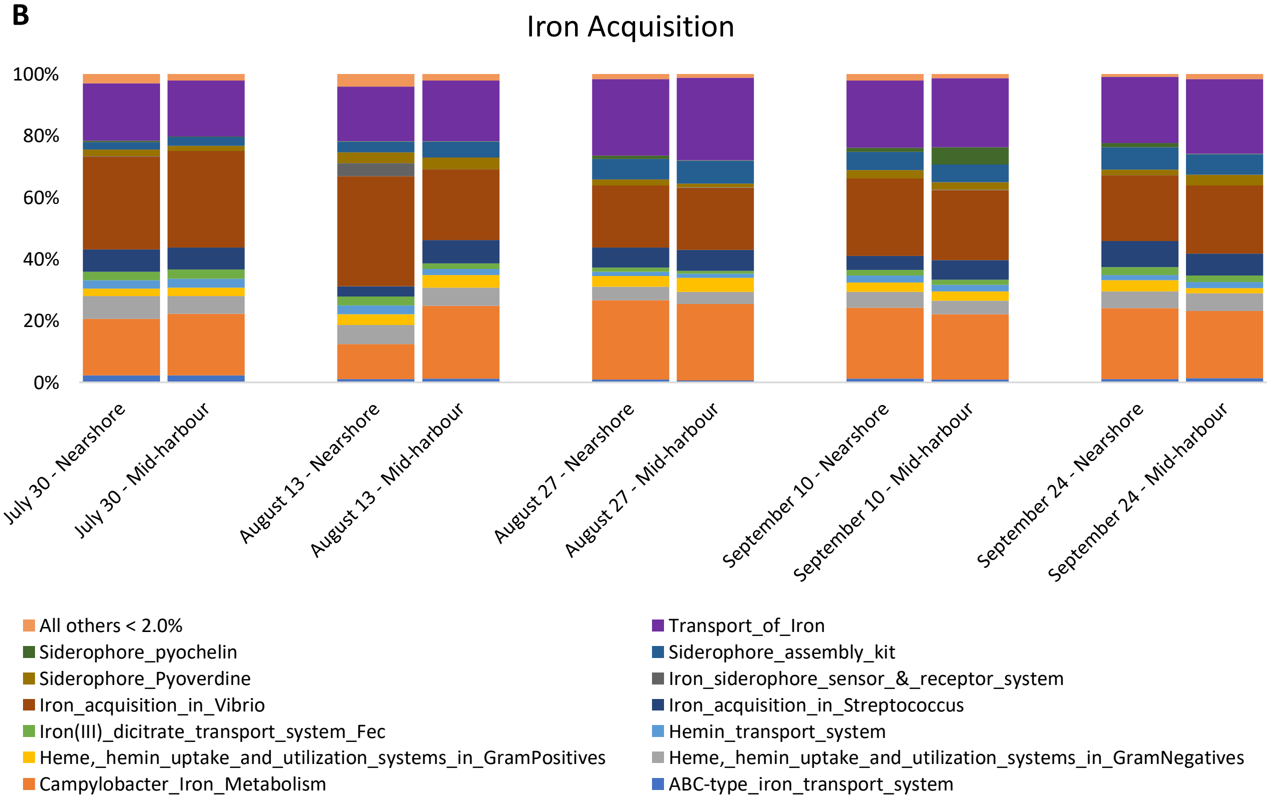
Iron Acquisition
100%
80%
60%
40%
20%
0%
July 30 - Nearshore
July 30 - Mid-harbour
August 13 - Nearshore
August 13 - Mid-harbour
August 27 - Nearshore
August 27 - Mid-harbour
September 10 - Nearshore
September 10 - Mid-harbour
September 24 - Nearshore
September 24 - Mid-harbour
All others < 2.0%
Transport_of_Iron
Siderophore_pyochelin
Siderophore_assembly_kit
Siderophore_Pyoverdine
Iron_siderophore_sensor_&_receptor_system
Iron_acquisition_in_Vibrio
Iron_acquisition_in_Streptococcus
Iron(III)_dicitrate_transport_system_Fec
Hemin_transport_system
Heme,_hemin_uptake_and_utilization_systems_in_GramPositives
Heme,_hemin_uptake_and_utilization_systems_in_GramNegatives
Campylobacter_Iron_Metabolism
ABC-type_iron_transport_system

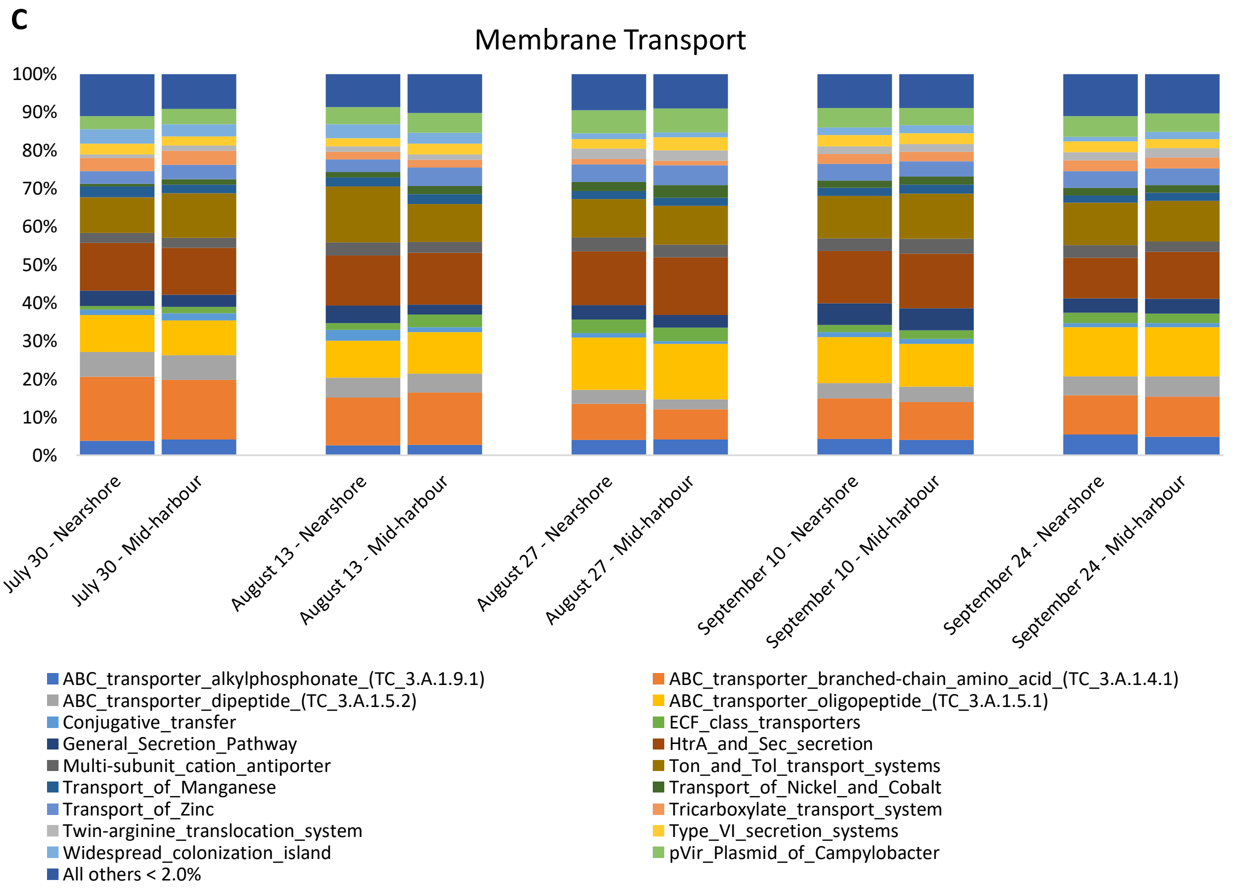


**Supplementary Figure 1.** Relative abundances of SEED level 1 functional categories comprising genes encoding A) potassium metabolism, B) iron acquisition, and C) membrane transport. In each panel, the relative abundance shown is the percent of the total number of assigned reads within each grouping. Only subcategories comprising > 5.0% of the total number genes in each category are included.
