## Supplementary Figure 2 for "Analysis of prokaryotic genes in an impaired Great Lakes harbour reveals seasonal metabolic shifts and a previously undetected cyanobacterium"

**A**

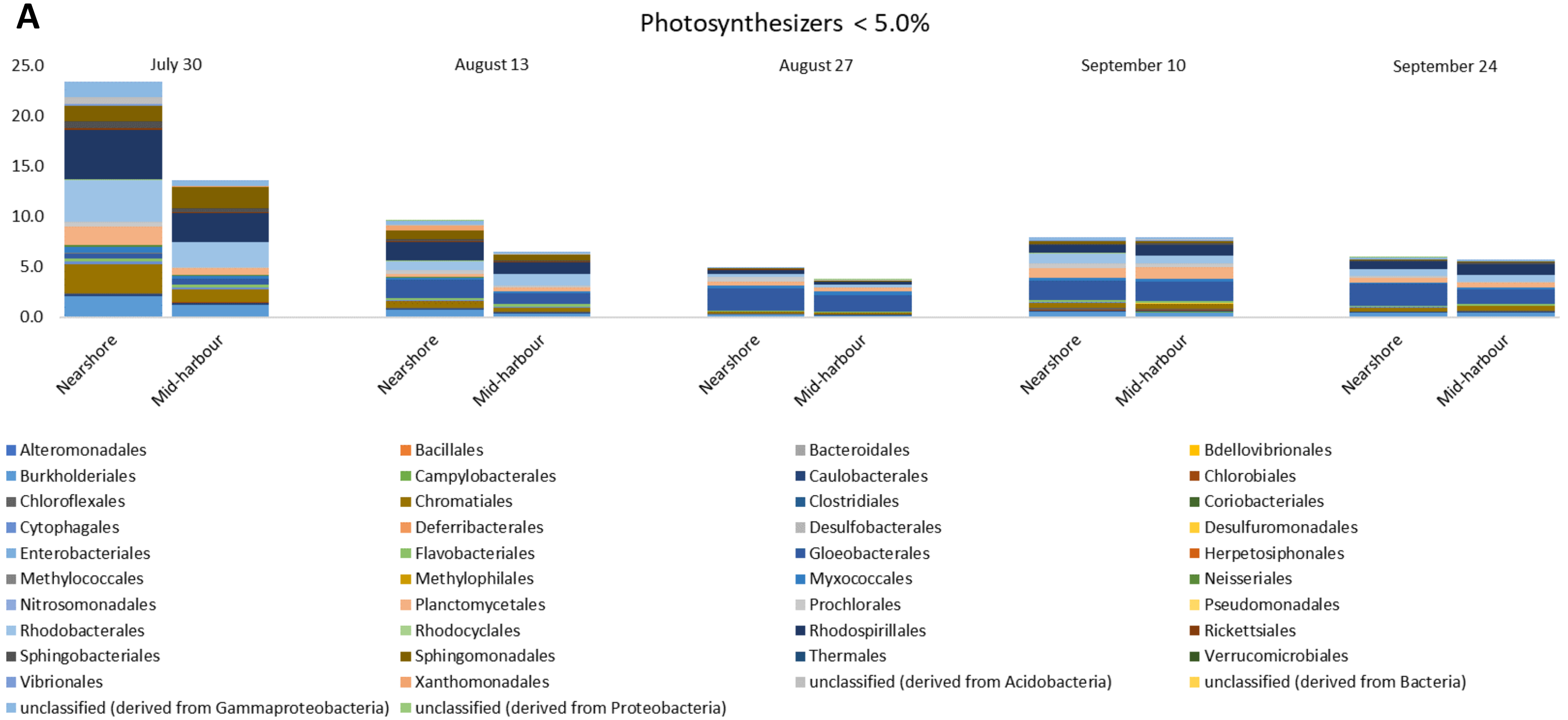
Photosynthesizers < 5.0%
July 30
August 13
August 27
September 10
September 24
25.0
20.0
15.0
10.0
5.0
0.0
Nearshore
Mid-harbour
Alteromonadales
Bacillales
Bacteroidales
Bdellovibrionales
Burkholderiales
Campylobacterales
Caulobacterales
Chlorobiales
Chloroflexales
Chromatiales
Clostridiales
Coriobacteriales
Cytophagales
Deferribacterales
Desulfobacterales
Desulfuromonadales
Enterobacteriales
Flavobacteriales
Gloeobacterales
Herpetosiphonales
Methylococcales
Methylophilales
Myxococcales
Neisseriales
Nitrosomonadales
Planctomycetales
Prochlorales
Pseudomonadales
Rhodobacterales
Rhodocyclales
Rhodospirillales
Rickettsiales
Sphingobacteriales
Sphingomonadales
Thermales
Verrucomicrobiales
Vibrionales
Xanthomonadales
unclassified (derived from Acidobacteria)
unclassified (derived from Bacteria)
unclassified (derived from Gammaproteobacteria)
unclassified (derived from Proteobacteria)

1

**B**

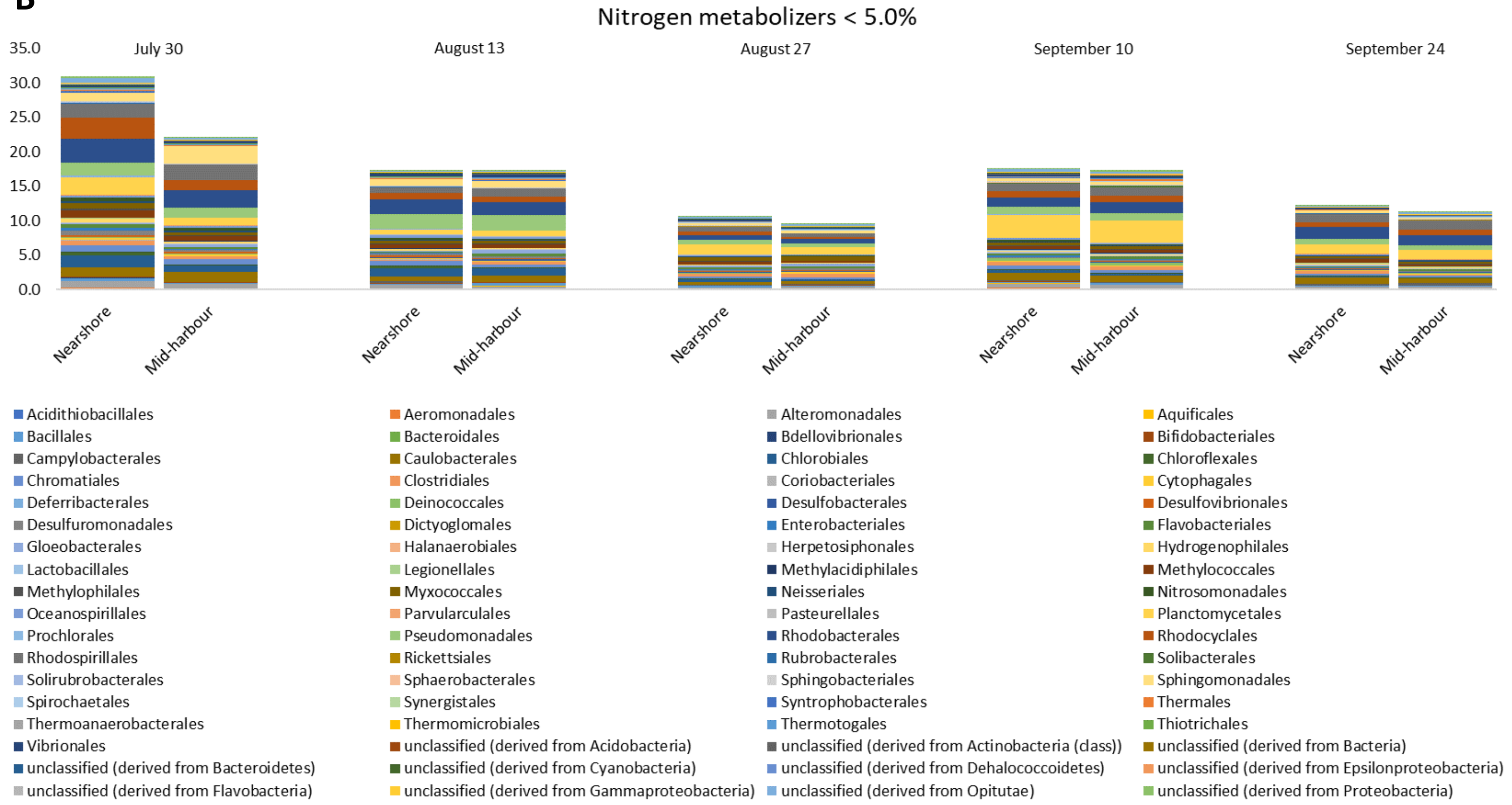
Nitrogen metabolizers < 5.0%
35.0
30.0
25.0
20.0
15.0
10.0
5.0
0.0
July 30
August 13
August 27
September 10
September 24
Nearshore
Mid-harbour
Nearshore
Mid-harbour
Nearshore
Mid-harbour
Nearshore
Mid-harbour
Nearshore
Mid-harbour
Acidithiobacillales
Aeromonadales
Alteromonadales
Aquificales
Bacillales
Bacteroidales
Bdellovibrionales
Bifidobacteriales
Campylobacterales
Caulobacterales
Chlorobiales
Chloroflexales
Chromatiales
Clostridiales
Coriobacteriales
Cytophagales
Deferribacterales
Deinococcales
Desulfobacterales
Desulfovibrionales
Desulfuromonadales
Dictyoglomales
Enterobacteriales
Flavobacteriales
Gloeobacterales
Halanaerobiales
Herpetosiphonales
Hydrogenophilales
Lactobacillales
Legionellales
Methylacidiphilales
Methylococcales
Methylophilales
Myxococcales
Neisseriales
Nitrosomonadales
Oceanospirillales
Parvularculales
Pasteurellales
Planctomycetales
Prochlorales
Pseudomonadales
Rhodobacterales
Rhodocyclales
Rhodospirillales
Rickettsiales
Rubrobacterales
Solibacterales
Solirubrobacterales
Sphaerobacterales
Sphingobacteriales
Sphingomonadales
Spirochaetales
Synergistales
Syntrophobacterales
Thermales
Thermoanaerobacterales
Thermomicrobiales
Thermotogales
Thiotrichales
Vibrionales
unclassified (derived from Acidobacteria)
unclassified (derived from Actinobacteria (class))
unclassified (derived from Bacteria)
unclassified (derived from Bacteroidetes)
unclassified (derived from Cyanobacteria)
unclassified (derived from Dehalococcoidetes)
unclassified (derived from Epsilonproteobacteria)
unclassified (derived from Flavobacteria)
unclassified (derived from Gammaproteobacteria)
unclassified (derived from Opitutae)
unclassified (derived from Proteobacteria)

2

C

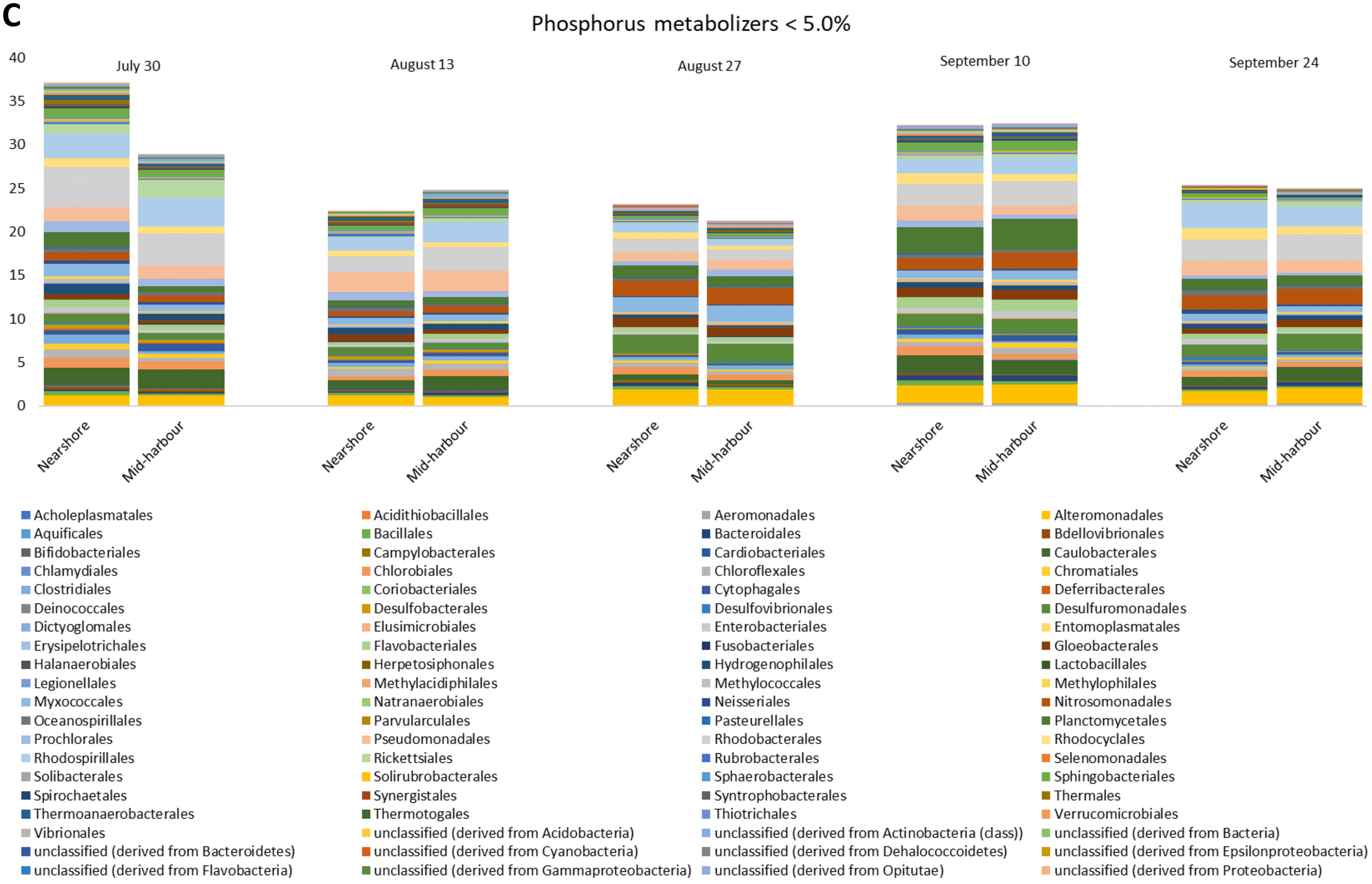
Phosphorus metabolizers < 5.0%
40
35
30
25
20
15
10
5
0
July 30
August 13
August 27
September 10
September 24
Nearshore
Mid-harbour
Acholeplasmatales
Acidithiobacillales
Aeromonadales
Alteromonadales
Aquificales
Bacillales
Bacteroidales
Bdellovibrionales
Bifidobacteriales
Campylobacterales
Cardiobacteriales
Caulobacterales
Chlamydiales
Chlorobiales
Chloroflexales
Chromatiales
Clostridiales
Coriobacteriales
Cytophagales
Deferribacterales
Deinococcales
Desulfobacterales
Desulfovibrionales
Desulfuromonadales
Dictyoglomales
Elusimicrobiales
Enterobacteriales
Entomoplasmatales
Erysipelotrichales
Flavobacteriales
Fusobacteriales
Gloeobacterales
Halanaerobiales
Herpetosiphonales
Hydrogenophilales
Lactobacillales
Legionellales
Methylacidiphilales
Methylococcales
Methylophilales
Myxococcales
Natranaerobiales
Neisseriales
Nitrosomonadales
Oceanospirillales
Parvularculales
Pasteurellales
Planctomycetales
Prochlorales
Pseudomonadales
Rhodobacterales
Rhodocyclales
Rhodospirillales
Rickettsiales
Rubrobacterales
Selenomonadales
Solibacterales
Solirubrobacterales
Sphaerobacterales
Sphingobacteriales
Spirochaetales
Synergistales
Syntrophobacterales
Thermales
Thermoanaerobacterales
Thermotogales
Thiotrichales
Verrucomicrobiales
Vibrionales
unclassified (derived from Acidobacteria)
unclassified (derived from Actinobacteria (class))
unclassified (derived from Bacteria)
unclassified (derived from Bacteroidetes)
unclassified (derived from Cyanobacteria)
unclassified (derived from Dehalococcoidetes)
unclassified (derived from Epsilonproteobacteria)
unclassified (derived from Flavobacteria)
unclassified (derived from Gammaproteobacteria)
unclassified (derived from Opitutae)
unclassified (derived from Proteobacteria)

3

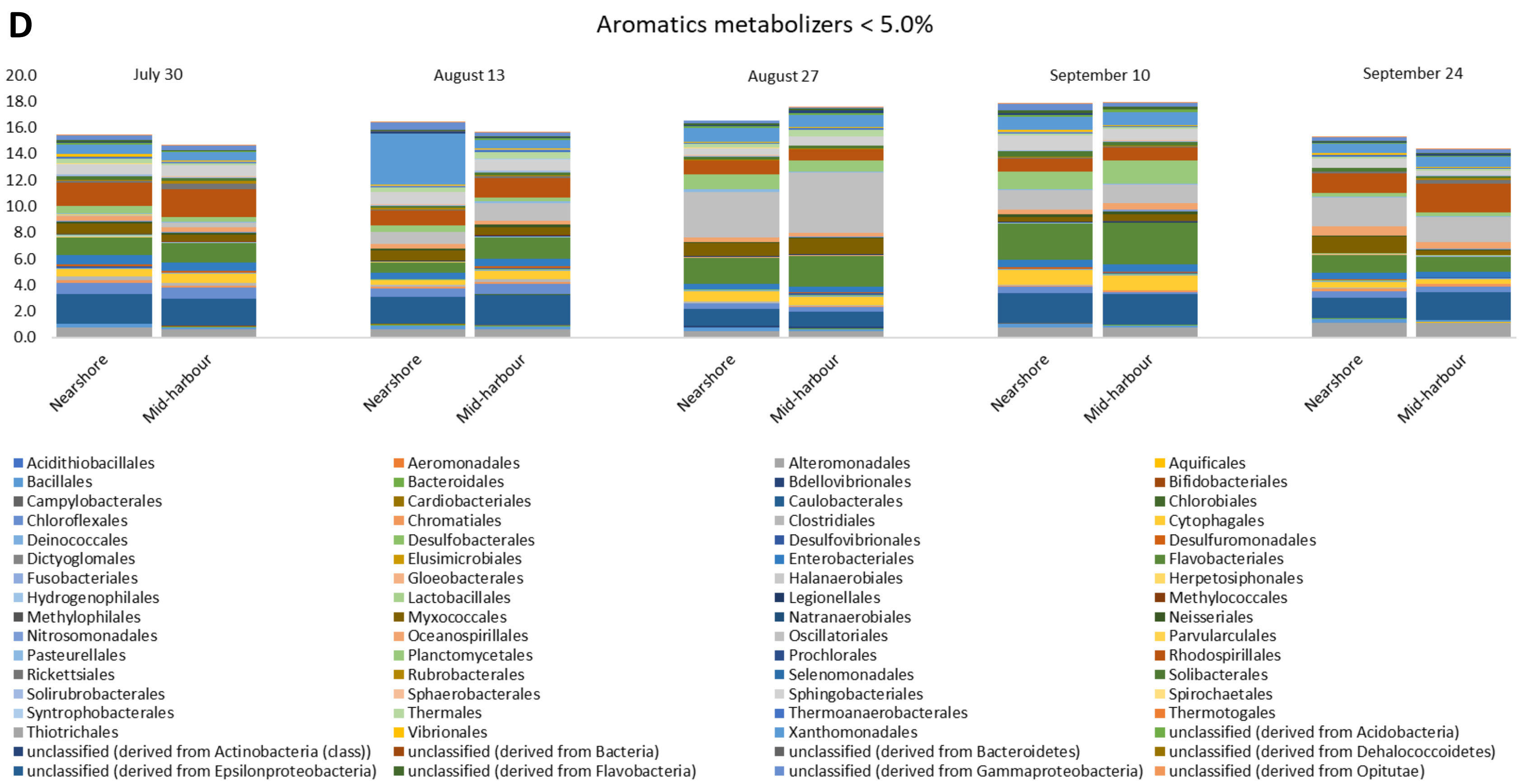


**Supplementary Figure 2.** Relative abundances of bacterial orders comprising < 5.0% of the community encoding genes for A) photosynthesis, B) nitrogen metabolism, C) phosphorus metabolism, and D) aromatic compound metabolism. In each panel, the relative abundance shown is the percent of the total number of assigned reads within each grouping.
